# Interplay of RAP2 GTPase and the cytoskeleton in Hippo pathway regulation

**DOI:** 10.1101/2023.10.10.561687

**Authors:** Chenzhou Wu, Xiaomin Cai, Ying Wang, Carlos D. Rodriguez, Lydia Herrmann, Giorgia Zoaldi, Chun-Yuh Huang, Xiaoqiong Wang, Viraj R. Sanghvi, Rongze O. Lu, Zhipeng Meng

## Abstract

The Hippo signaling is instrumental in regulating organ size, regeneration, and carcinogenesis. The cytoskeleton emerges as a primary Hippo signaling modulator. Its structural alterations in response to environmental and intrinsic stimuli control Hippo kinase cascade activity. However, the precise mechanisms underlying the cytoskeleton regulation of Hippo signaling are not fully understood. RAP2 GTPase is known to mediate the mechanoresponses of Hippo signaling *via* activating the core Hippo kinases LATS1/2 through MAP4Ks and MST1/2. Here we show the pivotal role of the reciprocal regulation between RAP2 GTPase and the cytoskeleton in Hippo signaling. RAP2 deletion undermines the responses of the Hippo pathway to external cues tied to RhoA GTPase inhibition and actin cytoskeleton remodeling, such as energy stress and serum deprivtion. Notably, RhoA inhibitors and actin disruptors fail to activate LATS1/2 effectively in RAP2-deficient cells. RNA sequencing highlighted differential regulation of both actin and microtubule networks by RAP2 gene deletion. Consistently, Taxol, a microtubule-stabilizing agent, was less effective in activating LATS1/2 and inhibiting cell growth in RAP2 and MAP4K4/6/7 knockout cells. In summary, our findings position RAP2 as a central integrator of cytoskeletal signals for Hippo signaling, which offers new avenues for understanding Hippo regulation and therapeutic interventions in Hippo-impaired cancers.

## 1. Introduction

The Hippo signaling pathway has consistently emerged as a central player in the orchestration of organ size, tissue regeneration, and carcinogenesis [1,2]. Its core comprises a kinase cascade including MST1/2-MAP4K and LATS1/2, which have the critical function of phosphorylating and thereby inactivating YAP and TAZ, two transcriptional co-factors vital for transcribing genes for cell proliferation, morphogenesis, and mobility [3]. These phosphorylation events play an instrumental role in maintaining cellular homeostasis and ensuring regulated tissue growth and development.

Recent advancements in the field have highlighted the significance of the cellular cytoskeleton as a modulator of the Hippo pathway [4-9]. Structural and mechanical alterations of the cytoskeleton, triggered by a range of environmental and intrinsic stimuli, have been proposed to directly and profoundly tune the core kinase cascade of the Hippo signaling pathway, specifically the activity of LATS1/2 [10]. Such changes can ripple through downstream processes, culminating in modifications to cell proliferation, apoptosis, and other vital cellular functions. However, despite these insights, the detailed mechanisms through which the actin and tubulin cytoskeletons regulate the Hippo pathway remain to be fully elucidated.

RAP2 GTPase, as recent investigations have established, is intricately tied to the mechanoresponses of Hippo signaling [11,12]. It exerts influence by activating the MAP4Ks and MST1/2 kinases while concurrently inhibiting the RhoA GTPase. These findings have spurred further interest in the precise role and regulation of RAP2 GTPase, especially considering its documented bidirectional relationship with the cytoskeleton [13-17] and its potential implications on Hippo pathway regulation. Several studies have documented the physical binding of the actin cytoskeleton to RAP2 GTPase [13,17], underscoring the importance of this interplay. Moreover, there is growing evidence suggesting that the actin cytoskeleton might wield influence over RAP2 activity, and conversely, RAP2 GTPase could modulate actin dynamics. This reciprocal regulation has far-reaching implications, especially in the context of how stress and biochemical cues regulate the Hippo pathway.

Our current study probes the intricate dynamics between RAP2 GTPase and the cytoskeleton in the contexts of stress and biochemical cues, seeking to shed light on the complex interplay between Hippo signaling and the cytoskeleton. We aim to provide a comprehensive understanding that could pave the way for therapeutic interventions in cancer malignancies and other diseases characterized by Hippo pathway disruptions.

## 2. Materials and Methods

### 2.1 Cell culture and treatment

HEK293A cells, a gift from Dr. Kun-Liang Guan’s group at UC San Diego, were cultured in DMEM supplemented with 10% fetal bovine serum. MCF10A cells were originally obtained from ATCC and cultured in DMEM/F12 supplemented with 5% horse serum, 20 ng/ml of human EGF, 0.5 mg/ml of hydrocortisone, 100 ng/ml of cholera toxin and 10 μg/ml of Insulin. The H-Ras V12 proto-oncogene transformed MCF10A cell line MCF10AT, a gift from Dr. Kun-Liang Guan, was also maintained in the above medium. For conditions of low cell density, we seeded 1.5 × 10^5^cells in each well of six-well plates. For a denser cell population, we either seeded 6.0 × 10^5^or 8 × 10cells in each well. For other treatments, cells at low density were exposed to 2-deoxy-D-Glucose (2-DG) at a concentration of 12.5 or 25 mM, Latrunculin B (LatB) at 0.075-0.5 μg/ml, Exoenzyme C3 at 1 μg/ml, Cytochalasin D (CytoD) at 100 nM, Lysophosphatidic acid (LPA) at 1 μM, Sorbitol at 200 mM, Isobutylmethylxanthine, 1-Methyl-3-Isobutylxanthine/Forskolin (IBMX/FSK) at 100 and 10 μM, respectively, and Paclitaxel (Taxol) at 3 μM.

For the purpose of inducing gene expression *via* tetracycline, we used the pRetroX-Tet-on-advanced and pRetrox-Tight-Puro plasmids from Clontech Inc. Stable HEK293A cell lines were developed following the guidelines provided by the manufacturer. To trigger gene expression of RAP2A, we administered 500 ng/ml of Doxycycline for a duration of at least 24 hours. MCF10AT cells were exposed to 50 nM of Taxol for cell viability and migration assays.

### 2.2. Generation of the Hippo component knockout cell lines

The CRISPR gene-editing method was employed to remove targeted genes. We cloned the guide RNA sequences into the px459 plasmids (Addgene 48319), kindly provided by Dr. Feng Zhang. These modified plasmids were then introduced into either HEK293A or MCF10A cells. A day post-transfection, cells that had been transfected were enriched using 1 μg/ml puromycin for 3 days. Following this, the cells were distributed into 96-well plates, ensuring just a single cell per well. The resulting clones were assessed via Western blot using gene-specific antibodies. We ensured that at least two distinct clones were employed for each gene deletion in the subsequent experiments. The guide RNA sequences for SAV1 and RAP2 from the previous studies were used [11,18]. All the information on single clones is listed as below:

RAP2 A/B/C tKO MCF10AT single clones, RAP2 A/B/C tKO HEK293A single clones and RAP2 A/B/C MST1/2 5KO HEK293A single clones were previously established [11]. MST1/2 KO, SAV1 KO and MAP4K4/6/7 tKO HEK293A single clones were previously established [19]. RAP2 A/B/C SAV1 4KO single clone was established from RAP2 A/B/C tKO single clone. MAP4K4/6/7 SAV1 4KO single clone was established from MAP4K4/6/7 tKO single clone. MAP4K4/6/7 RAP2 A/B/C 6KO was established from MAP4K4/6/7 tKO single clone. The KO efficiency, including SAV1 and RAP2, of these single clones was confirmed by western blot.

### 2.3. Biochemical assays

Western blot was performed following standard methods, mostly with 9-12& SDS PAGE gels. 7.5% phos-tag gel was used to resolve the phosphor-YAP proteins and was then transferred to PVDF membranes and probed with indicated antibodies. The information on the antibodies is provided in Supplementary Table 1. The GST-Ral-GDS pulldown for measuring RAP2 activity was performed as previously described [11].

### 2.4. Cell viability and migration assay

To test the impact of taxol on the cell viability of MCF10AT cells, cells were seeded in 96 well-plates at a density of 2.0 × 10^3 per well. After 24 hours, cells were treated with 50 nM of taxol for another 24 hours and tested. Cell viability was assessed by MTS using CellTiter 96® Aqueous One Solution Cell Proliferation Assay (Promega). For the transwell migration study, 5.0 × 10^4^cells were seeded in 12-well transwell inserts with 8 μm pore size. After 24 hours, the medium in the upper chamber was changed to 500 μl of no serum DMEM/F12, and 1ml of complete MCF10AT culture medium was added into the lower chamber. Taxol was only added in the upper chamber. After 24 hours, cells were fixed with 10% formalin and visualized with 0.1% crystal violet.

### 2.5. Bioinformatic analysis

The raw data from GSE98547 [11] was downloaded from the GEO database. Differential gene expression analysis was conducted using DESeq2 [20], with the criteria of log2FoldChange > 0 and a p-value < 0.05 to identify genes that exhibited differential expression. Gene Ontology (GO) Cellular Component analysis [21] was carried out in R (version 4.3.1), and the top 10 enriched terms were visualized in a plot.

## 3. Results

### 3.1. RAP2 mediates Hippo regulation by environmental signals and stresses

We previously reported that RAP2 GTPase mediates the Hippo pathway regulation by matrix stiffness by two downstream effectors, MAP4Ks and ARHGAP29 [11]. While the former is a direct kinase of LATS1/2, the later can activate the kinase cascade of MAP4K-MST1/2 and LATS1/2 through RhoA inhibition and actin depolymerization. Therefore, we speculated that RAP2 GTPases, including RAP2A, RAP2B, and RAP2C, may be also involved in the Hippo pathway regulation by other environmental biochemical signals and stresses beyond mechanical cues. Therefore, we challenged RAP2A/B/C triple knockout (RAP2A/B/C-tKO) cells and parental cells with various signaling activators or stress inducers.

G protein-coupled receptors (GPCRs) modulate the Hippo signaling pathway through the interactions of their downstream Gα proteins with the actin cytoskeleton and Rho GTPases [7,22]. Upon activation by specific ligands, GPCRs, depending on the Gα protein they couple to, promote polymerization or depolymerization of F-actin and subsequently tune down or up the activity of LATS1/2, respectively. Gαs-coupled GPCRs activate LATS1/2 kinases through the actin depolymerization by the cyclic AMP (cAMP)-PKA signaling, which was also observed in this study where the co-application of forskolin (FSK, an adenylyl cyclase activator elevating cAMP) and 3-Isobutyl-l-methylxanthine (IBMX, a phosphodiesterase inhibitor preventing cAMP breakdown) quickly induced phosphorylation of LATS at its hydrophobic motif (HM) and subsequent phosphorylation of YAP at Serine 127, a classical residue for LATS phosphorylation (**Fig. 1A**). In contrast, the phosphorylation of LATS and YAP was strongly compromised in the RAP2A/B/C-tKO cells.

**Figure 1.**
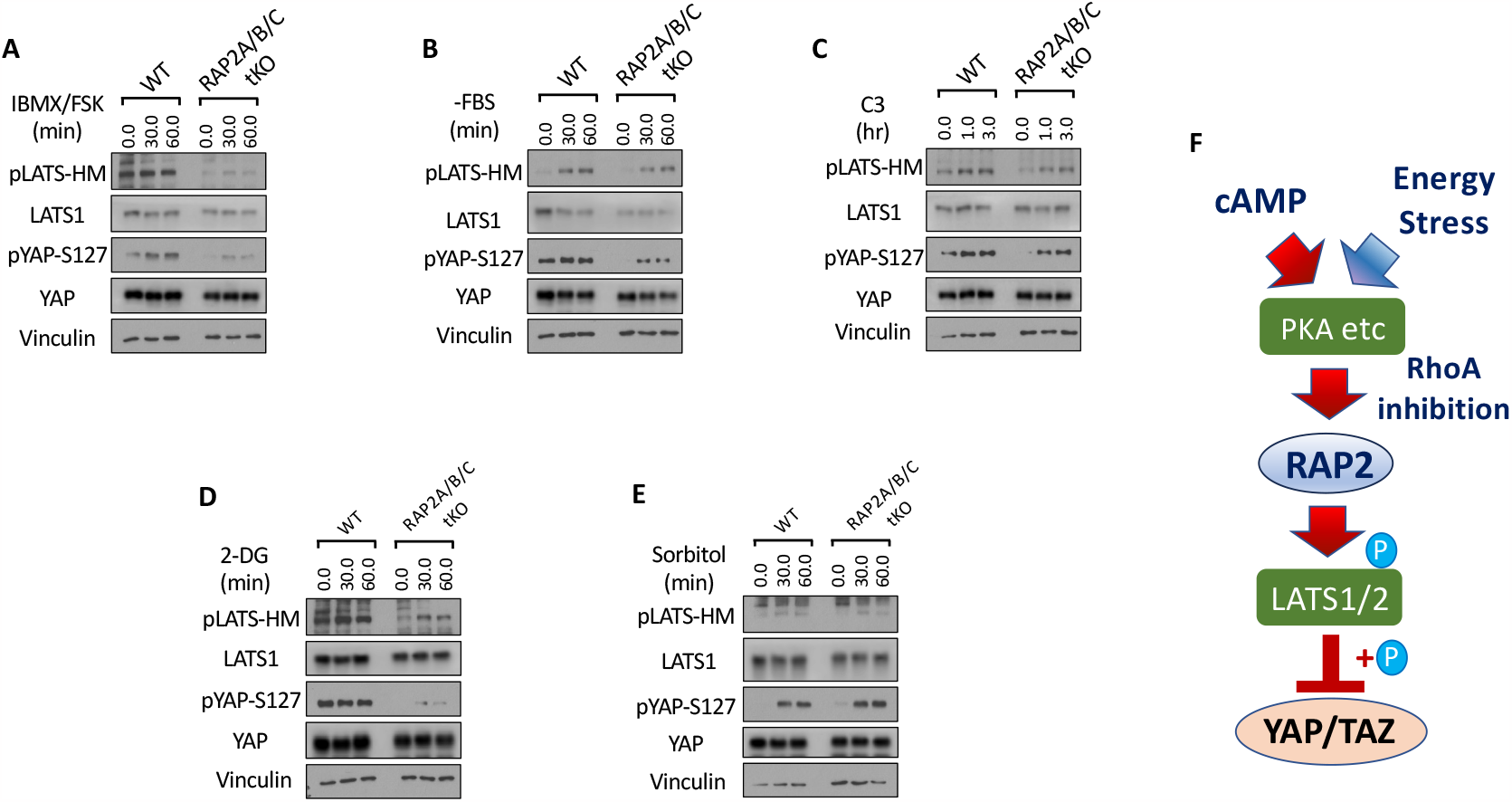
RAP2 mediates Hippo regulation by environmental signals and stresses. **A**. Wildtype (WT) and RAP2A/B/C triple knockout (tKO) cells were treated with cAMP inducers IBMX and FSK for 30 and 60 minutes. Then the cell lysates were collected for analyses of LATS phosphorylation at hydrophobic motif (HM) and YAP phosphorylation at Serine 127. **B**. WT and RAP2A/B/C tKO cells were serum starved, by replacing 10% FBS DMEM with serum-free DMEM (-FBS), for 30 and 60 minutes. **C**. WT and RAP2A/B/C tKO cells were treated with the Rho inhibitor C3 for 1.0 and 3.0 hours for analyses of the role of RAP2 in Hippo regulation by Rho GTPase. **D**. WT and RAP2A/B/C tKO cells were treated with an energy stress-inducer 2-deoxy-D-glucose (2-DG) for 30 and 60 minutes. **E**. WT and RAP2A/B/C tKO cells were treated with an energy stress-inducer sorbitol for 30 and 60 minutes. **F**. A diagram of a working model illustrating the role of RAP2 GTPase in the Hippo signaling regulation by environmental signals and stress.

The LATS phosphorylation and activation caused by increased cAMP is mainly atributed to RhoA inhibition and actin depolymerization mediated by PKA. On the other hand, the function of RAP2 GTPase is well-known to be regulated by direct actin binding. Considering the critical role of RhoA in Hippo signaling regulation documented in the previous literature, we first asked whether the RhoA inhibition acts through RAP2 GTPase to trigger LATS activation and YAP phosphorylation. Serum starvation has been reported to activate LATS by inactivating RhoA GTPase. Phosphorylation of YAP at Serine 127 was substantially compromised in the RAP2A/B/C tKO cells, though there is only a minor change in the phosphorylation of LATS hydrophobic motif (**Fig. 1B**). However, phosphorylation of LATS and YAP triggered by the treatment of the RhoA inhibitor the Exoenzyme C3 was also moderately compromised in the RAP2A/B/C tKO cells (**Fig. 1C**), indicating other proteins affected C3, other than those associated with actin cytoskeleton, could regulate the phosphorylation of LATS and YAP independently of RAP2 GTPase.

We further examined the role of RAP2 GTPase in Hippo signaling regulation by energy stress (**Fig. 1D**) and osmotic stress (**Fig. 1E**), respectively [23-25], to the RAP2A/B/C tKO and parental cells. The phosphorylation of LATS and YAP resulting from 2-deoxy-D-Glucose (2-DG) was greatly diminished in the RAP2A/B/C tKO cells, suggesting a prominent role of RAP2 GTPase in the regulation of Hippo signaling by energy stress (**Fig. 1D**). This result is consistent with a previous study showing that PKA knockout cells display weaker Hippo activation upon 2-DG-induced energy stress. In contrast, RAP2A/B/C tKO and parental cells responded similarly to hyperosmotic stress caused by sorbitol treatment (**Fig. 1E**). It should be noted that the role of RhoA in the Hippo signaling regulation by hyperosmotic stress has not been directly defined. These results altogether indicated that RAP2 GTPase is selectively engaged in the Hippo pathway regulation by certain extracellular signals.

In summary, the above results clearly showed that RAP2 GTPase can mediate the Hippo signaling regulation by environmental signals (i.e., Gs-coupled GPCR ligands and energy stress) other than matrix stiffness and other environmental cues (**Fig. 1F**). An interplay between RAP2 GTPase and RhoA GTPase was also implicated in the Hippo regulation in the contexts.

### 3.2. RAP2 is a required component of the MAP4K-mediated non-canonical Hippo signaling (*Hpo 2*) to respond to environmental signals

The MAP4K-mediated non-canonical Hippo signaling pathway is a molecular cascade that diverges from the classical Hippo pathway mediated by MST1/2. The biological roles of this MAP4K-Hippo signaling pathway have recently been implicated in the mouse models [26-28], and key Hippo components (SAV1, NF2, KIBRA) have thus been re-stratified (Hpo 1 vs. Hpo 2) [26].

Our previous study demonstrated that RAP2 acts through MAP4Ks to activate LATS1/2 during mechanosensing [11], positioning RAP2 as a component of the non-canonical MAP4K-Hippo signaling pathway (Hpo2)[26] (**Fig. 2A**). To test this notion, we applied different doses of 2-DG to induce differential levels of energy stress in MST1/2 dKO and RAP2A/B/C-MST1/2 5KO cells [11] in addition to RAP2A/B/C tKO cells. Though RAP2A/B/C deletion alone was already sufficient to block most of YAP hyperphosphorylation (seen by YAP band shift in phos-tag gel analysis), TAZ hyperphosphorylation (seen by TAZ band shift in regular SDS-PAGE) and LATS hydrophobic motif (HM) phosphorylation, combined deletion of RAP2A/B/C and MST1/2 nearly blocked all responses of YAP and LATS to energy stress (**Fig. 2B**), suggesting that RAP2 and MST1/2 work in parallel to activate LATS kinases in this context. The band shift of TAZ proteins, which indicates its hyperphosphorylation, also displayed a similar pattern.

**Figure 2.**
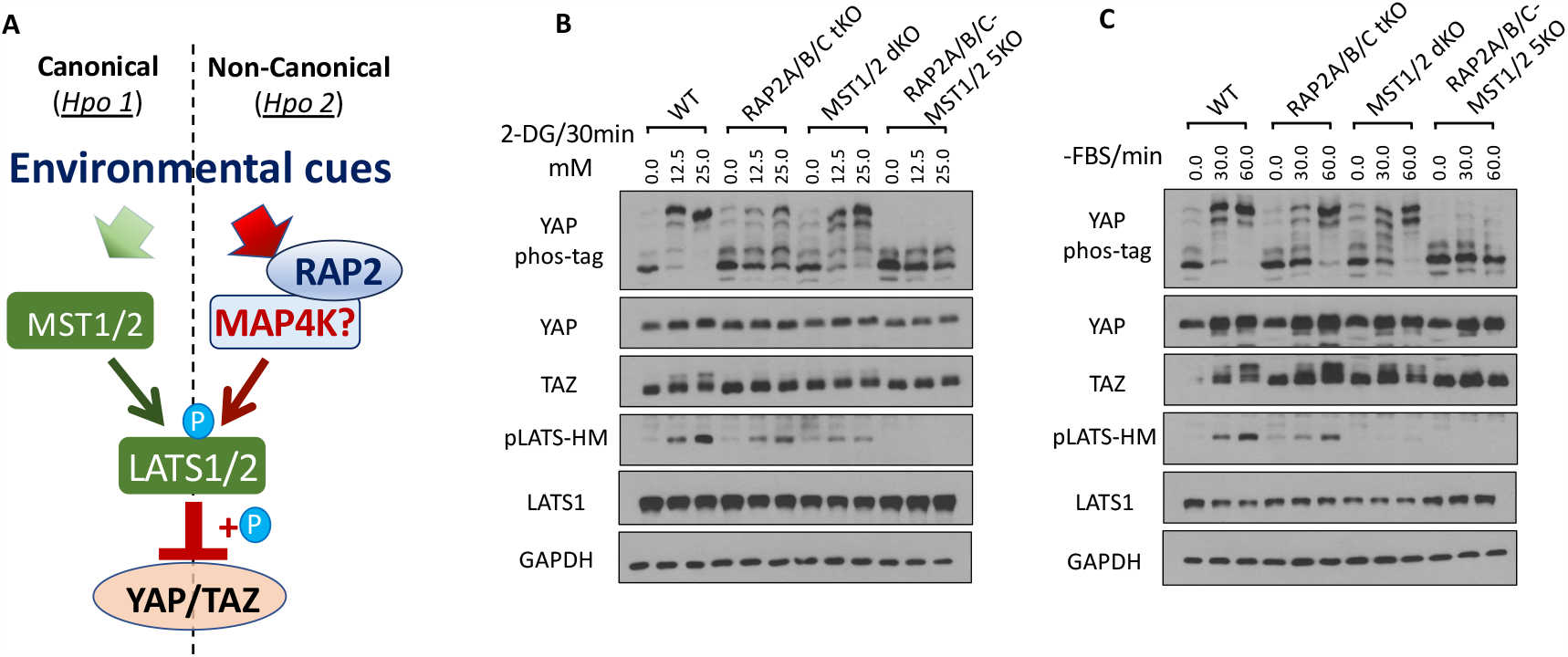
RAP2 is a required component of the MAP4K-mediated non-canonical Hippo signaling (*Hpo 2*) to respond to environmental signals. **A**. A diagram illustrating the role of RAP2 in the canonical MST1/2-mediated (Hpo1) and the non-canonical MAP4K-mediated (Hpo2) signaling pathways. **B**. WT, RAP2A/B/C tKO, MST1/2 dKO, and RAP2A/B/C-MST1/2 5KO cells were treated with low dose (12.5 mM) and high dose (25.0 mM) of 2-DG. **C**. WT, RAP2A/B/C tKO, MST1/2 dKO, and RAP2A/B/C-MST1/2 5KO cells were serum-starved for 30 and 60 minutes.

Furthermore, we applied serum starvation to deplete RhoA activity in the same set of knockout cells and discovered that the phosphorylation of LATS, YAP, and TAZ (shown by its band shift) could be blocked by deletion of both RAP2A/B/C and MST1/2, but not either alone (**Fig. 2C**). This result demonstrated that RAP2A/B/C and MST1/2 function as parallel components in mediating LATS1/2 activation by serum starvation.

Altogether, environmental signals can act through both RAP2A/B/C and MST1/2, likely in parallel, to activate LATS1/2 and inactivate YAP/TAZ, though the results did not rule out the crosstalk between RAP2A/B/C and MST1/2 as previously reported in mechanotransduction [11].

### 3.3. The Hippo signaling regulation by actin cytoskeleton requires RAP2 GTPase

RhoA GTPase controls Hippo pathway activity through modulating actin polymerization and stress fiber formation [3]. To understand whether RAP2 GTPase is required by Hippo activation by actin depolymerization resulting from RhoA inhibition, we applied Latrunculin B (LatB), which binds to actin monomers near the nucleotide-binding cleft and thus prevents actin polymerization and thus actomyosin contractile ability [29], to the set of RAP2A/B/C tKO, MST1/2 dKO, and RAP2A/B/C-MST1/2 5KO cells (**Fig. 3A**).

**Figure 3.**
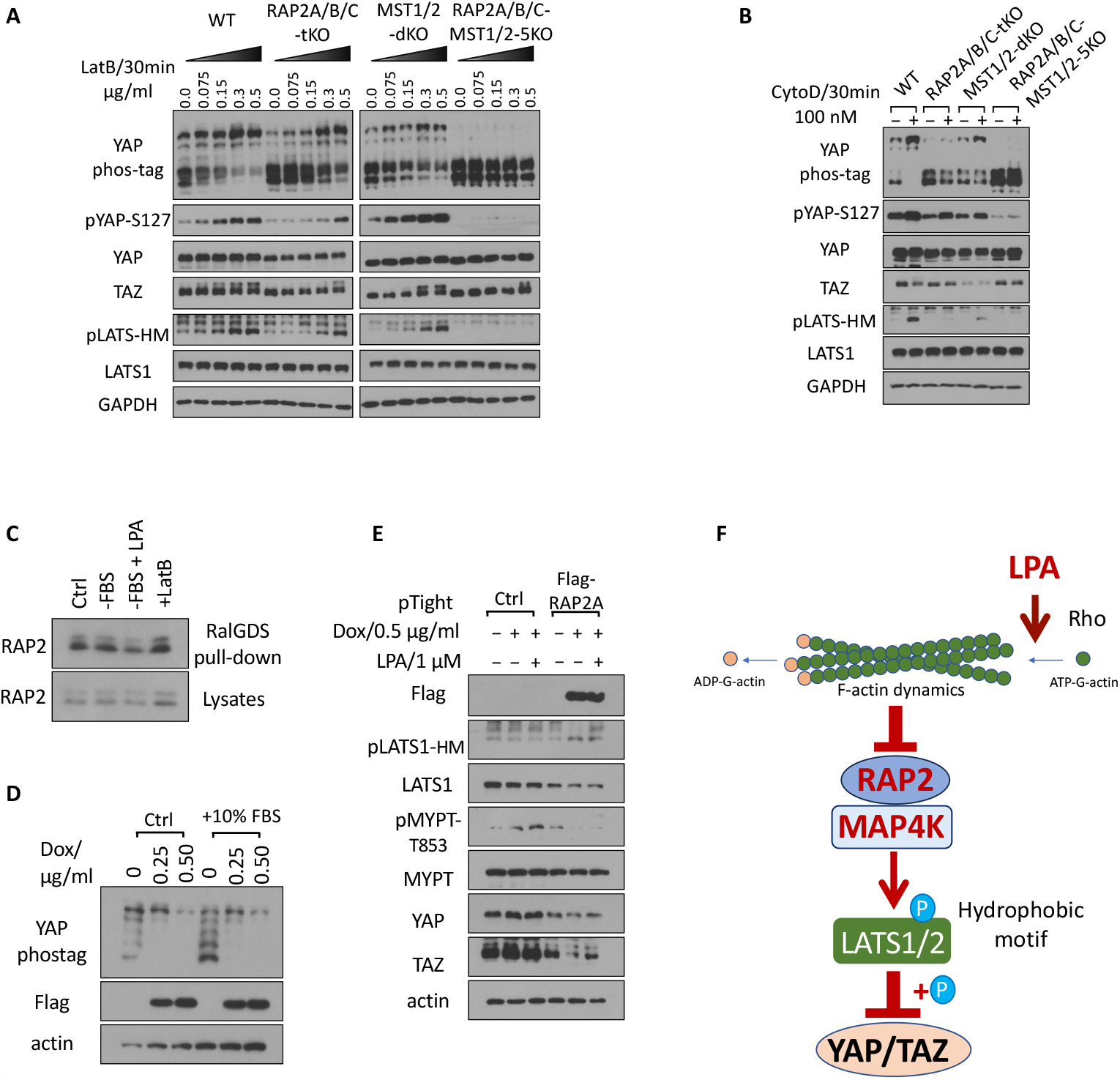
The Hippo signaling regulation by the actin cytoskeleton requires RAP2 GTPase. **A**. WT, RAP2A/B/C tKO, MST1/2 dKO, and RAP2A/B/C-MST1/2 5KO cells were treated with a range of doses of Latrunculin B (LatB) for 30 minutes. **B**. WT, RAP2A/B/C tKO, MST1/2 dKO, and RAP2A/B/C-MST1/2 5KO cells were treated with 100 nM Cytochalasin D for 30 minutes. **C**. Cells were serum starved for 1 hour (-FBS), or treated with 1 μM LPA after 1 hour serum starvation (-FBS + LPA), or treated with LatB for 30 minutes (+LatB). **D**. HEK293A cells transduced with pRetroX-Tet-on-advanced-neo viruses and selected by G418, and then were transduced with pRetroX-Tight-puro-Flag-RAP2A vectors and then selected by Puromycin. The subsequent stable cells were treated with 0.25 or 0.50 μg/ml Doxycycline (Dox) for 24 hours in 10% FBS and then another 10% FBS was added to cells. **E**. Cells transduced with the inducible Flag-RAP2A viral vector and the control vector were treated with 0.50 μg/ml Dox for 24 hours and then were treated with 1 μMLysophosphatidic Acid (LPA) for 30 minutes. **F**. A working model of how actin cytoskeleton dynamics acts through RAP2 GTPase to control the Hippo pathway activity.

We observed dose-dependent effects of LatB on the WT, RAP2A/B/C tKO, and MST1/2 dKO cells, but the effects of LatB in RAP2A/B/C tKO cells were greatly blunted. More importantly, deletion of both RAP2A/B/C and MST1/2 completely blocked the effect of LatB on YAP, TAZ, and LATS (**Fig. 3A**). We further confirmed the role of RAP2A/B/C in actin-mediated Hippo regulation by treating cells with Cytochalasin D (CytoD), which binds to the (+) end of F-actin to block the addition of new monomer actin subunits. We observed similar responses of the knockout cells to CytoD (**Fig. 3B**), though LatB and CytoD disrupt actin dynamics in different manners, further supporting the general role of RAP2 GTPase in mediating the Hippo pathway regulation by actin cytoskeleton.

Small GTPases regulate biological functions by directly binding to their downstream effectors [30]. Furthermore, previous reports showed that actin can bind to RAP2 GTPase regardless of its GTP/GDP binding status. We thus determined whether RhoA and actin cytoskeleton could regulate RAP2 GTPase activities with a classical RalGDS binding assay as previously described (**Fig. 3C**) [11]. Though serum starvation by itself only very slightly increases the binding of RAP2 to RalGDS, stimulating RhoA activity with lysophosphatidic acid (LPA) can decrease RAP2 binding to RalGDS while a high dose of LatB (0.5 μg/ml) can more substantially increase RAP2 binding to RalGDS.

To determine whether RAP2 acts downstream of RhoA, we generated a RAP2A-inducible expression cell line (**Fig. 3D**) Induction of RAP2 expression by doxycycline (Dox) triggered YAP phosphorylation. An additional 10% FBS, which can activate RhoA GTPase, can decrease YAP phosphorylation in the cells only when RAP2 expression is not induced (as shown in the 1vs. 4lane of the Phostag analysis) (**Fig. 3D**). We also applied a high dose of LPA (1 μM) to the inducible cell line, which showed that LPA failed to block LATS phosphorylation caused by RAP2A expression (**Fig. 3E**). Furthermore, phosphorylation of MYPT, which indicates actin stress fiber formation, was increased by LPA but strongly blocked by RAP2A induction. A similar expression change of TAZ, in response to LPA and RAP2 induction, was also observed.

Overall, these results established a signaling axis of external stimuli => Rho GTPase => actin remodeling => RAP2 GTPase => Hippo kinase cascade (**Fig. 3F**).

### 3.4. RAP2 and SAV1 work in parallel to promote YAP phosphorylation by LATS via MAP4K and MST1/2, respectively

Salvador homolog 1 (SAV1) is an essential component of canonical Hippo kinase cascade mediated by MST1/2. SAV1 forms a complex with MST1/2 and thus promotes autophosphorylation of MST1/2. Recent advances have also shown that SAV1 functions as a scaffold protein that anchors MST1/2 at the plasma membrane for the subsequent phosphorylation and activation of LATS1/2, as well as attenuates MST1/2 inactivation by the STRIPAK-PP2A complex. Furthermore, we have found that the LATS phosphorylation by CytoD and LatB is blunted in the SAV1 KO cells [18], indicating that SAV1 is involved in cytoskeleton regulation of LATS1/2 (**Fig. 4A**). Consistently, SAV1 KO also displayed reduced LATS phosphorylation caused by high confluence (**Fig. 4B**), though YAP phosphorylation was less affected by SAV1 deletion.

**Figure 4.**
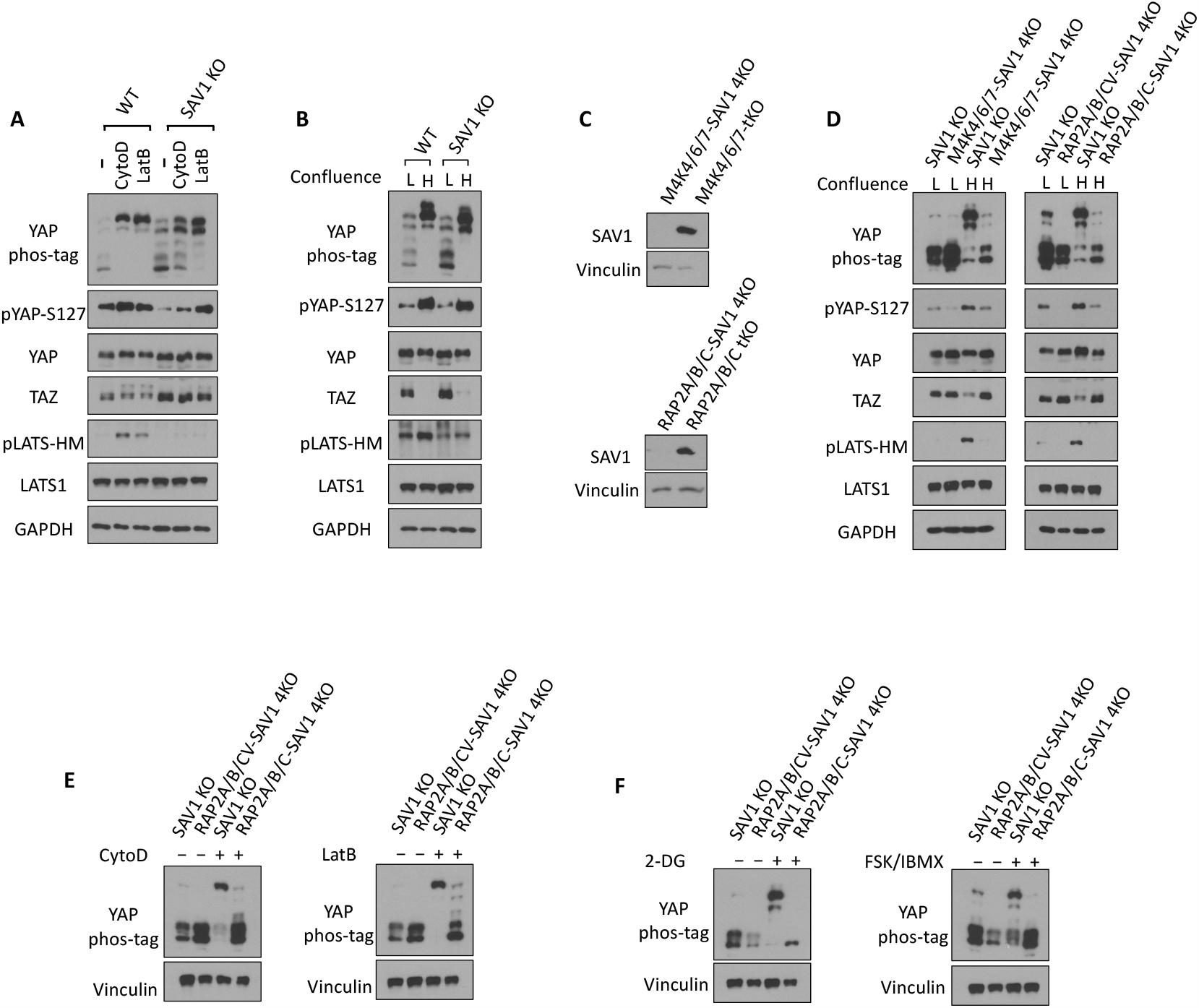
RAP2 and SAV1 work in parallel to promote YAP phosphorylation by LATS *via* MAP4K and MST1/2, respectively. **A**. WT and SAV1 KO cells were treated with 100 nM CytoD and 0.2 μg/ml LatB for 30 minutes. **B**. WT and SAV1 KO cells were grown at low and high (4.5x dense) confluence for 24 hours and then collected for immunoblot analysis. **C**. Validating the deletion of SAV1 in MAP4K4/6/7 tKO and RAP2A/B/C tKO cells by CRISPR/Cas9. **D**. SAV1 KO, MAP4K4/6/7-SAV1 4KO, and RAP2A/B/C-SAV1 4KO cells were grown at low and high confluence. **E**. SAV1 KO and RAP2A/B/C-SAV1 4KO cells were treated with 100 nM CytoD and 0.2 μg/ml LatB for 30 minutes. **F**. SAV1 KO and RAP2A/B/C-SAV1 4KO cells were treated 2-DG or FSK/IBMX for 30 minutes.

To understand the potential interaction and cooperation of SAV1 and the RAP2-MAP4K non-canonical Hippo signaling, we deleted SAV1 in MAP4K4/6/7 tKO and RAP2A/B/C tKO cells to generate MAP4K4/6/7-SAV1 4KO and RAP2A/B/C-SAV1 4KO cells (**Fig. 4C**). These two 4KO cell lines showed significantly compromised phosphorylation of YAP and LATS caused by high (H) confluence when compared with low (L) confluence (**Fig. 4D**), mimicking the responses of MAP4K4/6/7-MST1/2 5KO and RAP2A/B/C-MST1/2 5KO that were generated and characterized previously.

Similarly, phosphorylation of YAP in MAP4K4/6/7-SAV1 4KO and RAP2A/B/C-SAV1 4KO cells cannot be stimulated efficiently by actin disassembly-inducers CytoD and LatB (**Fig. 4E**), energy stress-inducer 2-DG, or PKA-activators FSK/IBMX (**Fig. 4F**). These results suggested the non-reductant and parallel functions of SAV1 and RAP2A/B/C or MAP4K4/6/7 in mediating Hippo pathway regulation by external signals, while RAP2A/B/C requires MAP4K4/6/7, but not the MST1/2 and SAV1 complex, to activate LATS1/2. To further support this notion, we deleted RAP2A/B/C gene in MAP4K4/6/7 tKO to examine whether it can potentiate Hippo deficiency in MAP4K4/6/7 tKO cells (**Fig. 5A**). In fact, RAP2A/B/C-MAP4K4/6/7 6KO cells and MAP4K4/6/7 6KO cells showed similar YAP hyperphosphorylation upon the treatment of LatB and serum starvation (**Fig. 5B, C**), though there was a different response to 2-DG-induced energy stress (**Fig. 5D**). Still, YAP hyperphosphorylation by 2-DG in RAP2A/B/C-MAP4K4/6/7 6KO cells cannot be blocked as much in RAP2A/B/C-MST1/2 5KO cells. These results altogether supported a working model where RAP2 functions upstream of MAP4K in the non-canonical Hippo (Hpo2) signaling [26] to mediate phosphorylation of LATS and YAP triggered by actin disassembly and stress (**Fig. 5E)**. The role of RAP2 in the Non-Canonical (Hpo2) signaling is as crucial as the well-characterized role of SAV1 in the Canonical (Hpo1) signaling.

**Figure 5.**
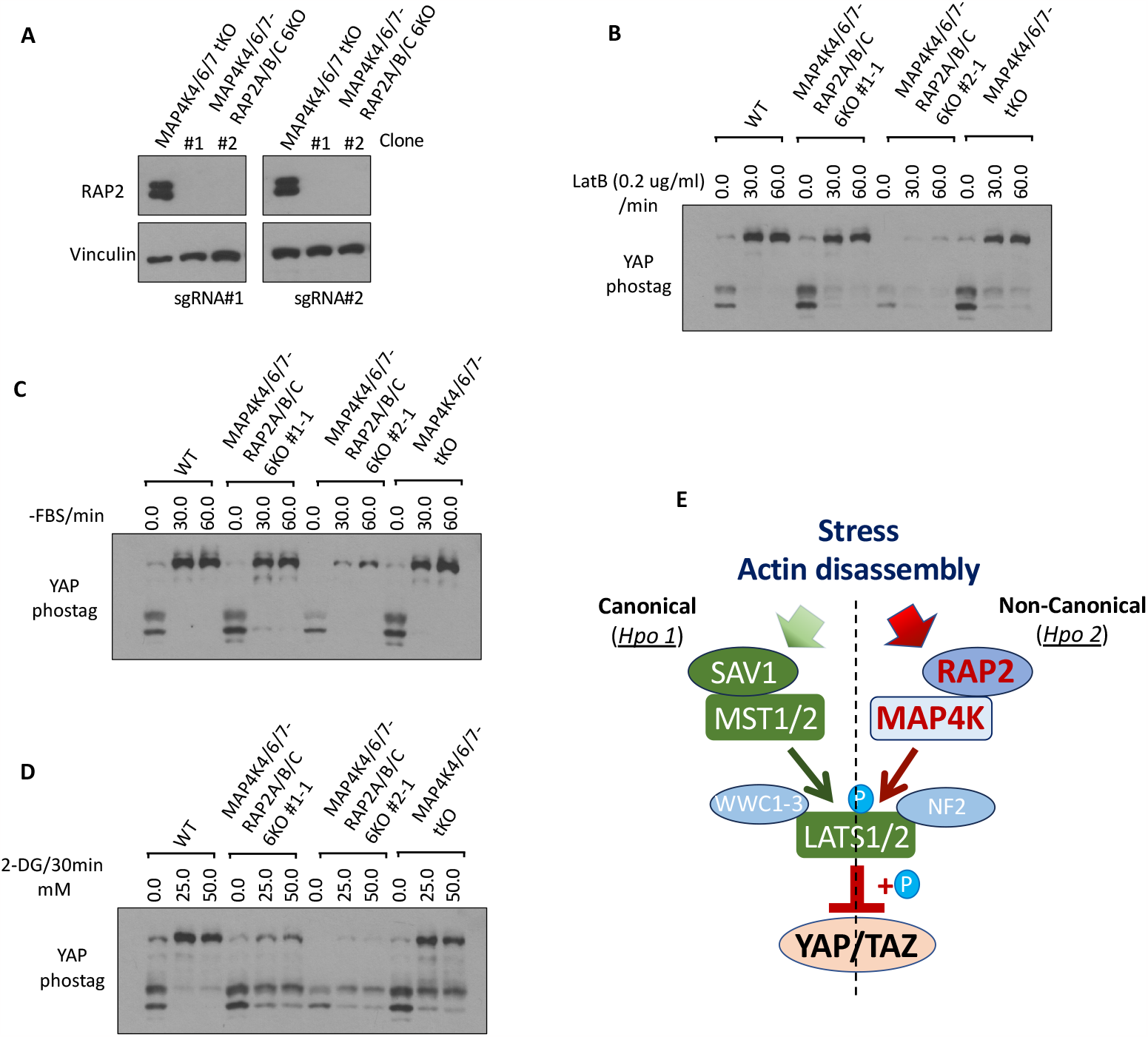
Combined deletion of RAP2 GTPase and MAP4Ks does not further potentiate Hippo deficiency compared with deletion of RAP2 GTPase or MAP4Ks alone. **A**. Two different sets of sgRNAs targeting RAP2A, RAP2B, and RAP2C genes, as previously described in Ref 11, were used to delete these 3 genes in MAP4K4/6/7 tKO cells. Two clones were expanded for further analysis. **B-D**. WT, MAP4K4/6/7 tKO, and RAP2A/B/C-MAP4K4/6/7 6KO cells were treated with LatB (**B**), serum starvation (**C**), and 2-DG (**D**) for 30 minutes. **E**. A diagram to show the parallel role of SAV1 and RAP2 GTPase in mediating signaling transduction of Hpo1 and Hpo2.

### 3.5. Interplay between RAP2 GTPase and cytoskeleton dynamics

Our gene ontology analyses of RNA sequencing of RAP2A/B/C tKO and WT cells at low stiffness (1 kPa) and high stiffness (40 kPa) (GSE98547) revealed the expression of many genes involved in actin cytoskeleton had been altered by RAP2A/B/C gene deletion, particularly at low stiffness where RAP2 GTPase was more active. In addition, at high stiffness, microtubule genes were also strongly differentially expressed in RAP2A/B/C tKO cells (**Fig. 6A**).

**Figure 6.**
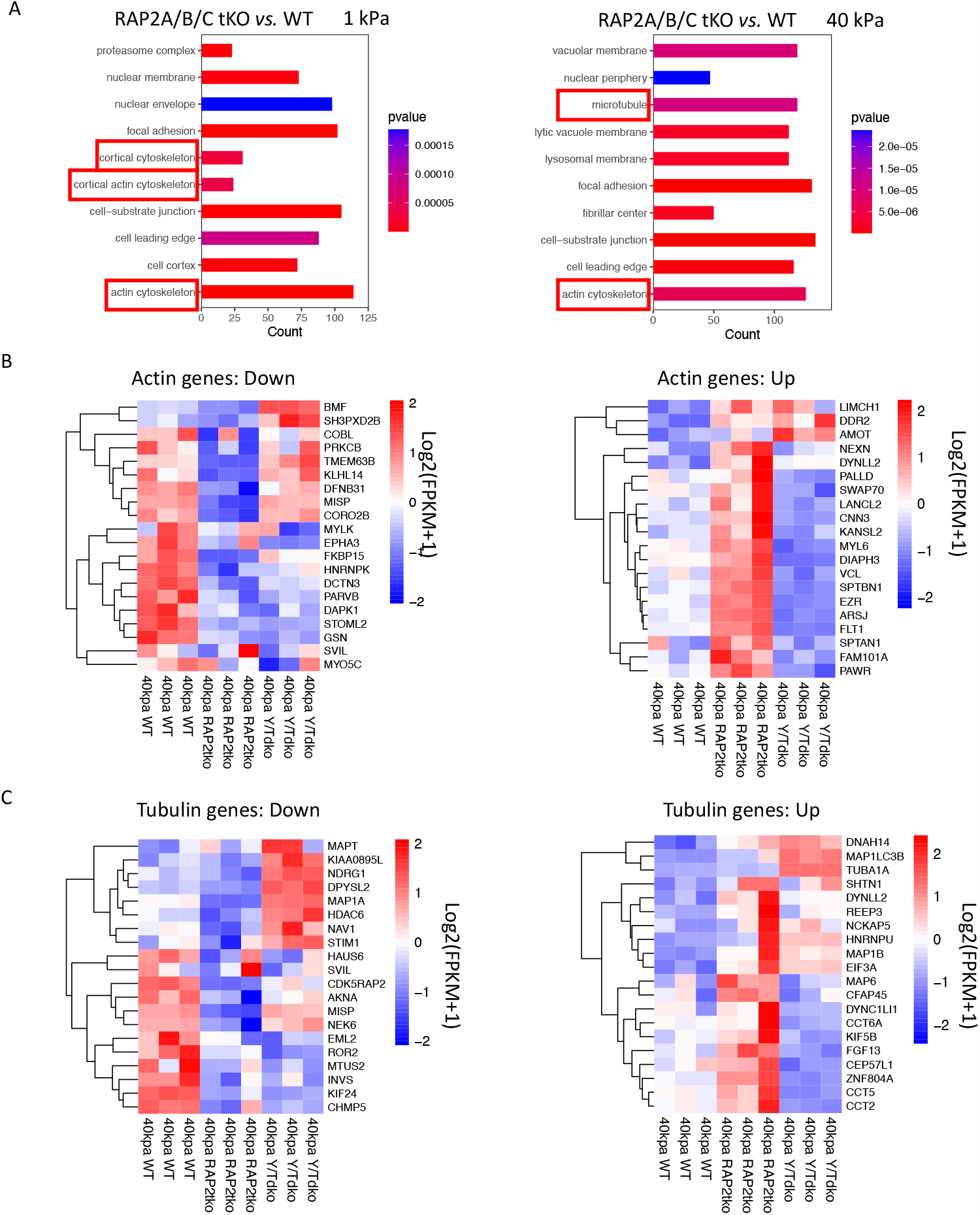
Differentially expressed actin and tubulin genes in RAP2A/B/C tKO cells. **A**. Gene ontology for cellular components with the RNAseq analysis of RAP2A/B/C tKO cells versus WT cells on sod (1 kPa) and stiff (40 kPa) substrates. Cytoskeleton terms were enriched. **B**. Down- and up-regulated actin-related genes, in WT, RAP2A/B/C tKO, and YAP/TAZ dKO (Y/T dKO) were ploted with heat-map. **C**. Down- and up-regulated tubulin-related genes, in WT, RAP2A/B/C tKO, and Y/T dKO were ploted with heat-map.

Many altered actin genes in RAP2/B/C tKO cells were potentially YAP/TAZ-targeting genes, as YAP/TAZ dKO (Y/TdKO) cells showed oppositely regulated patterns when compared with RAP2/B/C tKO cells (**Fig. 6B**), especially those up-regulated by RAP2A/B/C deletion at high stiffness. Many tubulin genes are also differentially regulated by RAP2 and YAP/TAZ (**Fig. 6B**), including microtubule-associated proteins (e.g., MAP1A, MAP6)[31] and chaperonins for tubulin and actin protein folding (e.g., CCT2, CCT5, CCT6A)[32].

As RAP2 appears to regulate tubulin dynamics in the cells, we asked whether RAP2 is involved in the Hippo pathway regulation by tubulin dynamics. Stabilization of tubulin by Taxol increases phosphorylation of YAP and LATS (**Fig. 7A**), which is consistent with the observation that nocodazole, a tubulin-disrupting agent, decreases phosphorylation of YAP and LATS. Interestingly, the effect of Taxol on YAP and LATS is reduced in RAP2A/B/C tKO and MAP4K4/6/7 tKO cells, suggesting that the RAP2-MAP4K signaling also plays a role in Hippo pathway regulation by tubulin dynamics. As Taxol has been often used as a first-line chemotherapy drug [33] and YAP is known to confer chemoresistance, we compared the responses of RAP2A/B/C tKO MCF10A cells that were transformed by the H-Ras V12 proto-oncogene (MCF10AT) [11] (**Fig. 7B**). In fact, the growth inhibition of Taxol on MCF10AT cells was substantially reduced by the deletion of RAP2A/B/C (**Fig. 7C**). Furthermore, although taxol strongly inhibits the migration of MCF10AT cells, this inhibition was also reduced in RAP2A/B/C tKO MCF10AT cells [11] (**Fig. 7D, E**). While a previous study implicated that MAP4K mediates the crosstalk of F-actin and microtubule [31], our study hereby suggested that the RAP2-MAP4K non-canonical Hippo signaling and the tubulin/actin cytoskeleton mutually regulate each other to control cell growth and mobility (**Fig. 5E**).

**Figure 7.**
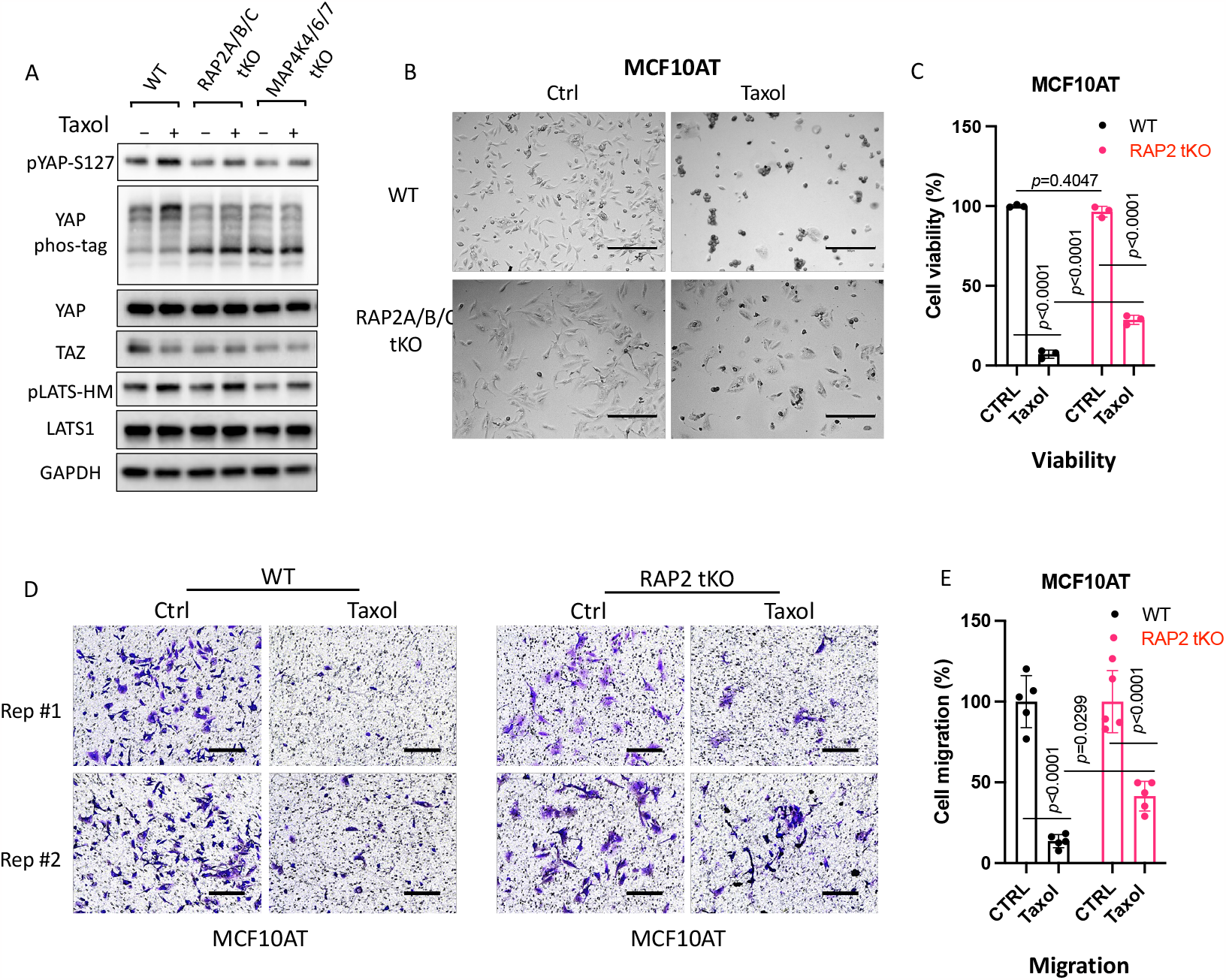
RAP2 GTPase is involved in Hippo signaling regulation by tubulin cytoskeleton dynamics. **A**. WT, RAP2A/B/C tKO, and MAP4K4/6/7 tKO HEK293A cells were treated with Taxol (3 μM) for 120 minutes. **B & C**. Representative bright filed images (**B**) and cell viability MTS test (**C**) showed cell survival of WT and RAP2A/B/C tKO MCF10AT cells treated with 50 nM of Taxol for 24 hours. For the MTS assay, a Two-way ANOVA and Tukey’s multiple comparison test was applied. ns, not significant; ****, p<0.0001. Scale bar, 300 μm. **D**. Transwell migration assay on WT and RAP2A/B/C tKO MCF10AT cells treated with 50 nM of Taxol for 24 hours. Scale bar, 100 μm. **E**. Quantification of cell migration assay. Student t-test was performed to determine the p value.

## 4. Discussion

### 4.1. RAP2 is an essential component of the non-canonical Hippo signaling pathway for sensing environmental and intrinsic cues

Understanding the mechanism by which organs perceive their size and regulate growth has been a major challenge in cellular biology. Central to this conundrum is the Hippo signaling pathway, an evolutionarily conserved network governing organ size, tissue regeneration, and tumorigenesis. In our study, we elucidate a novel aspect of this regulation. This study, together with previous studies [11,12], has implicated a crucial role of RAP2 GTPases in the Hippo pathway regulation by environmental signals, particularly those cytoskeleton-remodeling signals.

RAP2, much like NF2 and WWC1-3 [26], appears pivotal in facilitating and modulating kinase interactions within the Hippo pathway. Specifically, our results showcase RAP2’s capability to interface with LATS1/2 and upstream kinases such as MST1/2 and MAP4K1-7, providing a functional nexus for signal transduction.

Interestingly, our data seem to place RAP2 in an intricate regulatory network with WWC1-3 and NF2. The later two proteins have recently been posited as “loading docks” for LATS1/2, integrating upstream signals [26]. Drawing from our results, RAP2 might also function as a dynamic regulator, and RAP2 and SAV1 work synergistically to enhance the activation of Hpo2 (non-canonical, MAP4K-mediated) and Hpo1 (canonical, MST1/2-mediated) signaling, respectively. However, as RAP2 also regulates actin dynamics, our findings have not fully excluded the role of RAP2 in Hpo1 activation. For instance, in the energy stress context, the Hippo regulation by RAP2 likely engaged RAP2 downstream regulators other than MAP4Ks (**Fig. 5D**).

### 4.2. Interplay of RAP2 GTPase and the cytoskeleton

A major revelation from our study lies in understanding the intricate relationship between RAP2 GTPase and the cytoskeleton. It is well established that the cytoskeleton, with its vast network of filaments and tubules, orchestrates a myriad of cellular processes, including signaling cascades such as the Hippo pathway [3,10]. Our data suggests that RAP2, beyond its role in the Hippo pathway, might be integral in shaping cytoskeletal dynamics.

Previous studies have implicated the physical regulation of RAP2 GTPase by actin cytoskeleton [13-17]. Given the pivotal role of the cytoskeleton in mechanotransduction, the potential for RAP2 to act as a mechanosensor becomes an exciting avenue for future exploration.

Our findings highlight instances where RAP2, likely in its active form based on the previous study, modulates Hippo pathways in response to actin and tubulin dynamics. In the future, more characterization will be needed to determine the impact of RAP2 GTPase in the formation of stress fibers formation and alteration of cellular stiffness. Moreover, the migration patterns of these cells were notably different, suggesting a role for RAP2 in cell motility, possibly through cytoskeletal rearrangements. Given its newfound association with the cytoskeleton and the previous finding of RAP2’s role in sensing matrix stiffness, RAP2 can play a dual role: as a modulator of the Hippo pathway and a mediator of cellular responses to mechanical stimuli.

Furthermore, we noted potential functional interplays between RAP2 and known cytoskeleton-associated proteins from the RNA sequencing. Such interplays might be pivotal for cellular processes like adhesion, spreading, and even differentiation. It also raises the possibility of a feedback loop where RAP2-induced changes in the cytoskeleton could further influence Hippo pathway dynamics independent of its downstream effector MAP4K. To date, the detailed molecular mechanisms by which the Hippo pathway is regulated by the cytoskeleton still need to be defined. Identification of cytoskeleton-associated proteins connected to RAP2-mediated Hippo pathway regulation, which are missing in our current working models (Fig. 5E), will help seal this knowledge gap in the molecular mechanisms.

## 5. Conclusions

The interplay of Hippo signaling, RAP2 GTPase, and the cytoskeleton presents potential implications in diseases, especially those characterized by cytoskeletal abnormalities or altered mechanotransduction. Unraveling the exact mechanisms and interactions would not only provide a clearer picture of cellular signaling but also pave the way for therapeutic interventions targeting this interface.

## Author contributions

Conceptualization, Zhipeng Meng; Data curation, Chenzhou Wu, Xiaomin Cai, Giorgia Zoaldi; Formal analysis, Xiaomin Cai; Funding acquisition, Zhipeng Meng; Investigation, Chenzhou Wu, Ying Wang, Carlos Rodriguez, lydia Herrmann, Viraj Sanghvi and Rongze Lu; Methodology, Zhipeng Meng; Project administration, Zhipeng Meng; Resources, C.-Y. Charles Huang, Xiaoqiong Wang and Zhipeng Meng; Supervision, Zhipeng Meng; Writing – original draft, Chenzhou Wu and Zhipeng Meng; Writing – review & editing, C.-Y. Charles Huang, Xiaoqiong Wang, Viraj Sanghvi and Rongze Lu.

## Funding

**This research is mainly funded** by the National Institute of General Medical Sciences of the National Institutes of Health (NIH) under award number R35GM142504 to ZM.

ROL is supported by a grant from NINDS (R01NS126501). VRS is the recipient of the Department of Defense Concept Award (W81XWH2010549), Department of Defense Career Development Award (W81XWH2110377), Rally Foundation Independent Research Award (20IN31), Stanley J. Glaser Foundation Young Investigator Award (UM SJG 2022-24), NIH MIRA R35GM147497, and American Cancer Society’s Research Scholar Grant (RSG-22-071-01-TBE). ZM and VRS are both supported by faculty development and start-up funds from the University of Miami Miller School of Medicine and Sylvester Comprehensive Cancer Center. Research reported in this publication was supported by Sylvester Comprehensive Cancer Center under Translational Clinical Oncology Intra-Programmatic Award number TB-INTRA-2024-01 to ZM. LH was supported in part by an NCI R25 CREATE Summer Undergraduate Research Fellowship (R25CA261632).

## Conflicts of Interest

The authors declare no conflict of interest.

## Notes

### Competing Interest Statement

The authors have declared no competing interest.

